# Model-based species delimitation: are coalescent species reproductively isolated?

**DOI:** 10.1101/764092

**Authors:** Luke C. Campillo, Anthony J. Barley, Robert C. Thomson

## Abstract

A large and growing fraction of systematists define species as independently evolving lineages that may be recognized by analyzing the population genetic history of alleles sampled from individuals belonging to those species. This has motivated the development of increasingly sophisticated statistical models rooted in the multispecies coalescent process. Specifically, these models allow for simultaneous estimation of the number of species present in a sample of individuals and the phylogenetic history of those species using only DNA sequence data from independent loci. These methods hold extraordinary promise for increasing the efficiency of species discovery, but require extensive validation to ensure that they are accurate and precise. Whether the species identified by these methods correspond to the species that would be recognized by alternative species recognition criteria (such as measurements of reproductive isolation) is currently an open question, and a subject of vigorous debate. Here we perform an empirical test of these methods by making use of a classic model system in the history of speciation research, flies of the genus *Drosophila*. Specifically, we use the uniquely comprehensive data on reproductive isolation that is available for this system, along with DNA sequence data, to ask whether *Drosophila* species inferred under the multispecies coalescent model correspond to those recognized by many decades of speciation research. We found that coalescent based and reproductive isolation based methods of inferring species boundaries are concordant for 77% of the species pairs. We explore and discuss potential explanations for these discrepancies. We also found that the amount of prezygotic isolation between two species is a strong predictor of the posterior probability of species boundaries based on DNA sequence data, regardless of whether the species pairs are sympatrically or allopatrically distributed.

## INTRODUCTION

Due to the complex nature of the processes that drive speciation, the development of an operational species concept is difficult. For many years the Biological Species Concept (Mayr 1963), or some variation of it requiring high (but potentially incomplete) levels of reproductive isolation between populations, was the standard (Coyne and Orr 2004). Starting in the late 1990s, work echoing earlier evolutionary species concepts (Simpson 1951; Wiley 1978) led many evolutionary biologists to recognize that species might best be recognized as “metapopulation lineages” using any number of data types (de Queiroz 1998, 2007). Under this metapopulation conceptualization of species, the boundary between independent lineages (i.e., species) is demarcated by any number of criteria, which may or may not include reproductive isolation. Despite these conceptual changes, systematists remain in need of an operational framework for species delimitation. One option for placing the practice of species delimitation in a more explicit framework is to use statistical models of evolutionary relationships among genetic loci. One of the most widely used methods for doing so, BPP (Yang and Rannala 2014; Rannala and Yang 2017), is a Bayesian method based on the multispecies coalescent model (MSC). BPP can be employed to analyze variation in genealogies among many loci to identify independent coalescent lineages as species (Leaché and Fujita, 2010; Rannala and Yang, 2003).

These methods hold promise for greatly increasing the pace and accuracy of species description, although their utility depends on whether the lineages identified correspond to biologically defensible (or meaningful) species *and* the precision of these estimates. A number of studies have looked at the efficacy of the MSC for species tree inference (e.g., Fujita et al. 2012; Reid et al. 2014; Barley et al. 2018), but the concordance between independent coalescent lineages and species boundaries has been less well characterized in the literature. Relevant studies generally take one of two forms: 1) species boundaries identified under the MSC are compared to those identified by alternative approaches, with concordance viewed as an indication of accuracy (e.g., Camargo et al. 2012; Willis 2017); or 2) Data simulated under known demographic scenarios are analyzed under the MSC and the results are used to identify the relative over- or under-splitting of lineages by the model (Sukumaran and Knowles 2017; Barley et al. 2018; Luo et al. 2018). Both types of studies frequently find that the MSC identifies more lineages as species than some consider appropriate (Leaché et al. 2019). This discrepancy, compounded by the lack of a fully objective species criterion, has led to an increasing wariness that MSC delimitation may inflate the number of species, often beyond what many systematists find realistic.

These studies have been useful for characterizing the statistical performance of the MSC, but can be difficult to interpret in the context of real species radiations. On one hand, there is the possibility that the MSC identified genuine species that were missed by earlier studies because they are morphologically cryptic or otherwise difficult to differentiate. On the other hand, in simulation studies, the accuracy of results relies on the generating model (i.e., the simulation) being an accurate representation of biological reality. If important aspects of species biology are missing from the generating model we may draw inaccurate conclusions merely because our simulation study is overly simple relative to nature. The challenges associated with both of these approaches might be averted by turning to an empirical system where extensive data on the nature of species boundaries themselves are available. In such a system, we can examine the correspondence between lineages identified as being independent under the MSC and other criteria placed on species boundaries, such as reproductive isolation.

Here we make use of an empirical study system in which an extraordinarily rich understanding of reproductive isolation has been established for many species pairs, the genus *Drosophila*. We use this to compare species identification based on reproductive isolation with species delimitation under the MSC. *Drosophila* is an intensively studied model organism in many disciplines, and arguably the most influential clade for understanding the consequences of reproductive isolation as a mechanism of species formation and perpetuation. Much of this focus stems from the fact that experimental lab crosses are possible between species, which provides information on which species pairs show intrinsic reproductive isolation (lab crosses do not take extrinsic factors into account) and to what degree that isolation is complete. Two classic papers by Coyne and Orr (1989, 1997) synthesized data on reproductive isolation derived from experimental crosses and allozyme electrophoretic distance, which they used as a proxy for time since divergence by assuming a constant molecular clock. These data were used to produce estimates for the amount of total reproductive isolation (the combined accumulation of pre- and postzygotic isolation) and time (using genetic distance as a proxy) required for complete speciation. Here, complete speciation refers to a relaxed version of Mayr’s Biological Species Concept (Mayr 1963) that allows for low levels of gene flow between closely related species (see also Coyne and Orr 2004).

These studies provide insight into the nature of how species boundaries themselves become established. A central finding from Coyne and Orr (1989, 1997), and the one we focus on here, is that the amount of reproductive isolation increases with time since divergence. Building off this idea, we examined several aspects of how species identified under the MSC model matched those identified based on experimental quantification of reproductive isolation and genetic distance. Specifically, we were interested in measuring the correlation between the amount of reproductive isolation and genetic distance between two species, and the posterior probability that those two species were identified as independent coalescent lineages. By quantifying the relationship between reproductive isolation and coalescent lineages we stand to gain a better understanding of the relationship between the biological processes that drive speciation and promising statistical approaches to species delimitation. It should also be noted that most studies of reproductive isolation look at comparisons between species, whereas most practitioners of MSC species delimitation think about comparisons within species (i.e., at the population level). This study is explicitly concerned with delimitation of nominal species.

Assessing the accuracy of species delimitation remains difficult because speciation is a continuous process and species boundaries are not always discrete. However, utilizing information on reproductive isolation allows us to assess the performance of these methods in a more empirically grounded framework than is otherwise possible. With this in mind, the research goals for this study were to: 1) examine the relationship between coalescent and reproductive isolation based species delimitation; 2) quantify how varying levels of pre- and postzygotic isolation in allopatric and sympatric species pairs affect our ability to identify species under the MSC; 3) better understand the utility of MSC methods to increase the accuracy and precision of species recognition in the presence of partial, or complete, reproductive isolation.

## MATERIALS & METHODS

### Dataset Assembly

The lists of species pairs studied by Coyne and Orr (1989, 1997) were combined to assemble a set to be used in the present study (n = 108 pairwise comparisons). Values for pre- and postzygotic isolation were compiled from Coyne and Orr’s two studies (1989, 1997), and updated based on isolation measures from a more recent study (Yukilevich 2012). Total reproductive isolation (*T*; Coyne and Orr 1989) was calculated using the equation: *T* = *Pre* + (1 − *Pre*) *x Post*. In keeping with previous work, if a species pair only had data on prezygotic or postzygotic isolation, and that value was greater than 0.95 or 1.0, respectively, we considered *T* to equal the measure for which there was data (Coyne and Orr 1997). Conversely, if a species pair only had data for prezygotic or postzygotic isolation, and that value was less than 0.95 or 1.0, respectively, those species pairs were excluded from downstream analysis. Data on allozyme genetic distance (D) comes originally from Coyne and Orr (1989, 1997), but more recently updated by Yukilevich (2012).

For the purpose of this study, we follow the species recognition thresholds proposed by Coyne and Orr (1997). Specifically, they proposed minimum values on total reproductive isolation (T ≥ 0.903) and genetic distance (sympatric pairs: D ≥ 0.04; allopatric pairs: D ≥ 0.54) required for maintenance of species boundaries. While we recognize that the use of any particular threshold may raise concerns, we opted to use these for two reasons. First, by using the same metrics employed by previous studies, we are able to make a direct comparison between species delimitation based on reproductive isolation and delimitation based on MSC models. Second, because MSC species delimitation has many practical implications, we wanted to focus on values that would likely be the most relevant and widely used in empirical studies

We assembled a set of genes (n=8) that have been sequenced across a wide taxonomic breadth within *Drosophila* (van der Linde and Houle 2008; Yang et al. 2012). We then downloaded whatever sequence data was available from GenBank for this set of genes for all species (n = 59) on the compiled list. Custom Python and R scripts (R Core Team 2013) were used for data cleanup. Species pairs were then sorted into eleven species groups for analysis (Coyne and Orr 1997). Here, “species group” refers to a term used in *Drosophila* literature to reference a monophyletic set of closely-related species (see O’Grady and DeSalle 2018). While a large amount of mitochondrial data is available for *Drosophila*, we chose to include only one mitochondrial gene (Cytochrome c oxidase subunit II: COII) for each species group because we expect the entire mitochondrial genome to share a single coalescent history. We limited the final data matrix for each group to include only those genes that were missing sequence data for no more than two species. Because some intensively studied species have an abundance of sequence data relative to the other species in their group (e.g., *Drosophila subobscura* has 137 COII sequences available, while the other six species in the Obscura group combine for 30 available sequences), we pruned sequences from overrepresented taxa to make the amount of data more even across species. Specifically, for species that had >10 sequences at a given locus, we randomly pruned sequences until the number that remained was equal to the next most densely sampled species. Sequence data was aligned for each group using MUSCLE (Edgar 2004) and converted to PHYLIP format using DendroPy 4.2.0 (Sukumaran and Holder 2010). These eleven matrices, one for each species group, were analyzed independently in all downstream analyses. To check for any difference in electrophoretic distance and genetic distances calculated from DNA sequence data, we calculated Jukes-Cantor genetic distances across all species pairs for COII in Geneious v 7.1.9 (Kearse et al. 2012) and then compared those to Nei’s D (Fig. S1; Table S1).

### MSC Analysis

After assembling the dataset, we performed MSC species delimitation using BPP v 3.2 (Yang and Rannala 2014) on each of the 11 species groups separately. BPP uses reversible-jump Markov Chain Monte Carlo (rjMCMC) to estimate a Bayesian posterior distribution for different species delimitation models. We jointly estimated the species tree topology and assignment of individuals into species (referred to as “analysis A11” in BPP). Under this model we define a starting tree, where each named species in the group is a “population”, and then use the rjMCMC to target the posterior distribution of possible models that may merge combinations of these populations into a smaller number of species. This approach does not allow single populations to be split more finely, and so the existing taxonomy within each species group (i.e., named species from Coyne and Orr 1997) forms the upper limit on the total number of possible species in each analysis. We specified a prior distribution that assigns equal probability for the number of species (i.e., all populations merged into one species or a different species for each population) and then divides that probability by the proportion of compatible labeled histories (*speciesmodelprior = 2*; Yang 2015). We also set a prior that allows *Θ* (population size parameter) to vary among loci according to specified heredity multipliers (Hey and Nielsen 2004) to account for the differences in effective population size between nuclear and mitochondrial loci (*heredity = 2*; Yang and Rannala 2014).

Two central parameters of the MSC model are for population sizes (*Θ*) and species divergence times (*τ*). These are specified as gamma distributed random variables in BPP. Our goal was to set up a diffuse, but credible, prior distribution for both of these parameters. Here, we constructed an empirical prior for *Θ* in *Drosophila* by using published estimates of this value, if available, or published estimates of effective population size (*N*_*e*_) and mutation rate (*μ*) to solve for the equation: *Θ* = 4*Nμ* (Eyre-Walker et al. 2002; Wall et al. 2002; Tamura 2003; Yi et al. 2003; Hey and Nielsen 2004; Haag-Liautard et al. 2007; Pascual et al. 2007; Cutter 2008; Charlesworth 2009; Keightley et al. 2009, 2014; Legrand et al. 2009; Obbard et al. 2012; Smith et al. 2012; Schrider et al. 2013). Similarly, to obtain an empirical prior for the *τ* parameter, we used the online database TimeTree (http://www.timetree.org) to obtain an estimated root age for each species group. Because each species group had different estimated root ages, there was a unique combination of *Θ* and *τ* prior gamma distributions for each group. For both prior distributions, we centered the mean of the gamma distribution on the empirical values, and set the variance of the distribution to be wider than the distribution of all empirical values. We ran 204,000 MCMC generations, discarding the first 4000 generations as burnin and then sampling every 20 generations until we reached 10,000 samples (BPP output files will be available on Dryad Digital Repository upon publication). We repeated each analysis two times, checking that the results remained consistent across runs (Yang 2015).

The BPP manual (Yang 2015) explicitly states that there are no default priors for *Θ* and *τ*, yet many studies employ a set of three prior gamma distributions originally used by Leaché and Fujita (2010) as if they were. Here we explored prior sensitivity by performing additional BPP runs on each dataset under this suite of priors (Leaché and Fujita 2010). For the sake of clarity, we will refer to the set of priors from Leaché and Fujita (2010) as empirically “uninformed” and our empirically derived priors as “informed”. In total, we ran a BPP delimitation analysis on each of the eleven species groups under four unique prior settings (one informed, three uninformed) resulting in 44 independent BPP runs. Within the uninformed set of priors, we focus on those that most closely matched our prior expectations about the coalescent history for *Drosophila* (large ancestral population and shallow divergence among populations in this case). This should minimize differences between informed and uninformed priors, thereby supplying a realistic (and somewhat conservative) test of prior sensitivity. Full details for all prior settings can be found in supplementary material (Table S2), including *Θ* and *τ* priors for all groups and BPP runs. We extracted posterior *Θ* values from each MCMC output file using a custom Python script. To summarize the posterior distribution for *τ*, we used the R package Phytools v 0.6-60 (Revell 2012) to read the sampled trees in the MCMC output into R and calculate the tree height of each (Python and R codes will be available on Dryad Digital Repository upon publication). We visually inspected concordance between prior and posterior distributions of *Θ* and *τ* for the informed and uninformed priors. Additionally, we investigated whether the number of species delimited was sensitive to the alternative priors.

The A11 analysis in BPP estimates posterior probabilities for different delimitation models that may differ in number of species and the topology of the species phylogeny. Here, we are specifically interested in the marginal posterior probability for the splitting, or lumping, of each particular species pair, rather than the marginal posterior probability of any one delimitation model. Because species pairs can be split or lumped in different configurations, we calculated the marginal posterior probability of independence between each species in a given pair. We refer to this value as the Posterior Probability of Independent Lineages (PPIL). PPIL values are practically, and philosophically, the same measure as the “speciation probability” described in Leaché and Fujita (2010). The difference is that we calculate these values from analyses that marginalize across both species delimitations and species tree topologies, whereas Leaché and Fujita (2010) only marginalized across species delimitations due to limitations of the software at that time which required the species tree be held constant (i.e., we do not employ a guide tree in our analysis, as was done previously).

We calculated PPILs for all species pairs from Coyne and Orr (1989, 1997) where sufficient sequence data was available to conduct the analysis, and for which information on reproductive isolation was available (n = 108 pairwise comparisons. Using PPIL values, we compared the number of independent lineages identified by the MSC to those based on reproductive isolation and genetic distance documented by Coyne and Orr (1989, 1997). The two species in each pair were considered independent lineages if they had a high PPIL (≥ 0.95) or met the criteria for total isolation (T ≥ 0.903) and genetic distance (D_allo_ ≥ 0.54; D_symp_ ≥ 0.04) from Coyne and Orr (1997).

### Reproductive Isolation and Genetic Distance

Besides a cursory comparison in numbers of species delimited by either method, we were also interested in how the components of speciation considered by Coyne and Orr (1989, 1997) related to PPIL values in our present study. To accomplish this, we selected among a series of generalized additive models (GAMs) to assess the impact of different predictor variables (prezygotic isolation, postzygotic isolation, and genetic distance) on the PPIL for allopatrically and sympatrically distributed species pairs. We chose not to include total reproductive isolation in these models because it is a function of prezygotic and postzygotic isolation, which we already account for independently. We elected to use a GAM because the model can flexibly capture the impact of a predictor variable through a smoother function, allowing for both linear and non-linear relationships. Because the dependent variable (i.e., PPIL) included 0 and 1, we performed a logit transformation to normalize the data using the R package *car* (Fox and Weisberg 2011). We then used the R package *mgvc* to construct and fit GAMs to the data (Wood 2011). We calculated Akaike Information Criterion (Akaike 1974) values for each model and considered the model with the smallest AIC to be the best fitting model. We further considered models with ΔAIC ≤ 2 from the best model to be similarly plausible (Burnham et al. 2002).

We analyzed two independent sets of GAMs to: 1) explore the differences between allopatric and sympatric species pairs and 2) investigate the impact of predictor variables within allopatric and sympatric species pairs separately. The first series of GAMs compared models with one smoother function for all species pairs to models specifying separate smoothers for allopatric and sympatric species pairs. Improved fit for the latter model would indicate that range overlap between taxa differentially predicts PPIL values, and therefore these two types of species pairs (allopatric and sympatric) should be modelled separately in downstream analyses.

After confirming two smoothers based on geography was a better model, we fit the second series of GAMs on allopatric, sympatric, and closely related (D ≤ 0.50) allopatric and sympatric species pairs. For each set of species pairs, we fit a total of five GAMs, modelling the three predictor variables listed previously (prezygotic isolation, postzygotic isolation, and genetic distance) and two additive models (prezygotic isolation + genetic distance and postzygotic isolation + genetic distance). We fit the GAMs on taxa with low genetic distance (D ≤ 0.5) to maintain consistency with Coyne and Orr (1989), and because they suggested closely related sympatric species pairs should show the strongest signature of reinforcement. In this context, evidence for reinforcement’s effect would be that the model for prezygotic isolation is the strongest predictor of PPIL among closely related sympatric species, followed closely by a model of postzygotic isolation. Specifically, because reinforcement acts to increase prezygotic isolation in order to counteract disadvantageous hybridization, there must be some level of postzygotic isolation.

Because data from species pairs are phylogenetically non-independent, we also explored how this might impact our results. To do so, we extracted the maximum *a posteriori* tree from each of the BPP analyses using phytools v 0.6-60 (Revell 2012), setting branch lengths to their marginal posterior mean. We then assembled a reduced set of phylogenetically independent species pairs (i.e., only including pairs that are connected by paths on the tree that do not intersect with other pairs; see Felsenstein 2004, pg. 444). We then reran all the GAM analyses on this reduced but phylogenetically independent dataset.

Lastly, we investigated how PPIL values may vary with respect to levels of pre- and postzygotic isolation. To do this, we first constructed a reduced dataset that included all species pairs with prezygotic (n = 100) or postzygotic (n = 74) isolation data. As outlined by Coyne and Orr (1989, 1997), both of these values range from 0-1.0, with complete reproductive isolation (pre- or postzygotic) equal to 1.0. We then performed an ANOVA to determine if PPIL differed according to the levels of pre- and postzygotic isolation. Because the measures of postzygotic isolation increased from 0 to 1.0 in 0.25 increments, we treated each level as a separate category for the ANOVA. For prezygotic isolation, we binned the values based on the same levels (e.g., 0-0.25, 0.25-0.50, etc.). Coyne and Orr (1989, 1997) investigate what role Haldane’s Rule (Haldane 1922) may play in the speciation process and find it to be an important early step. We checked whether it might also influence PPIL by assembling a reduced dataset containing only those species pairs with 0.25-0.75 postzygotic isolation (n = 32), because at these levels of postzygotic isolation the pair could either be in line with Haldane’s Rule (i.e., the heterogametic sex, males in this case, are either sterile or infertile for both crosses) or not (i.e., a male and female, or both females are sterile/inviable). Information on sterility/infertility was taken from Yukilevich (2012). We manually categorized species pairs as either conforming to Haldane’s Rule or not and performed a one-way t-test to look for differences between both categories of species pairs.

## RESULTS

### Genetic dataset assembly and analysis

The final dataset included a total of 543 sequences from 8 genes. We were able to obtain sequence data for all species from one gene, COII, totaling 160 sequences for that gene. The median number of genes per species group was three. The Bipectinata group had the greatest gene diversity (n = 6), while the Buzzatii group was represented by only two genes. The mean number of sequences compared between each species pair in each species group was 7.25, ranging from an average of three sequences per pair in the Affinis group, up to 13.7 sequences per pair in the Melanogaster group. Complete information on number of genes per group and loci per species pair can be found in Supplementary Material (Table S3). The average genetic distance across all species pairs was 0.82, ranging from 0.026 (*D. heteroneura – D. silvestris*) to 1.95 (recorded for multiple species pairs in the Obscura group).

We ran the BPP analyses two times for each species group, and obtained similar results across runs. Consistency across runs indicates that mixing is likely adequate and the MCMC chain did not get stuck on any one model (Yang 2015). Species trees for each species group revealed varying levels of posterior probability support across nodes, but overall the species trees correspond with results from other phylogenetic studies (Fig. 1; van der Linde and Houle 2008; Yang et al. 2012).

**Figure 1).**
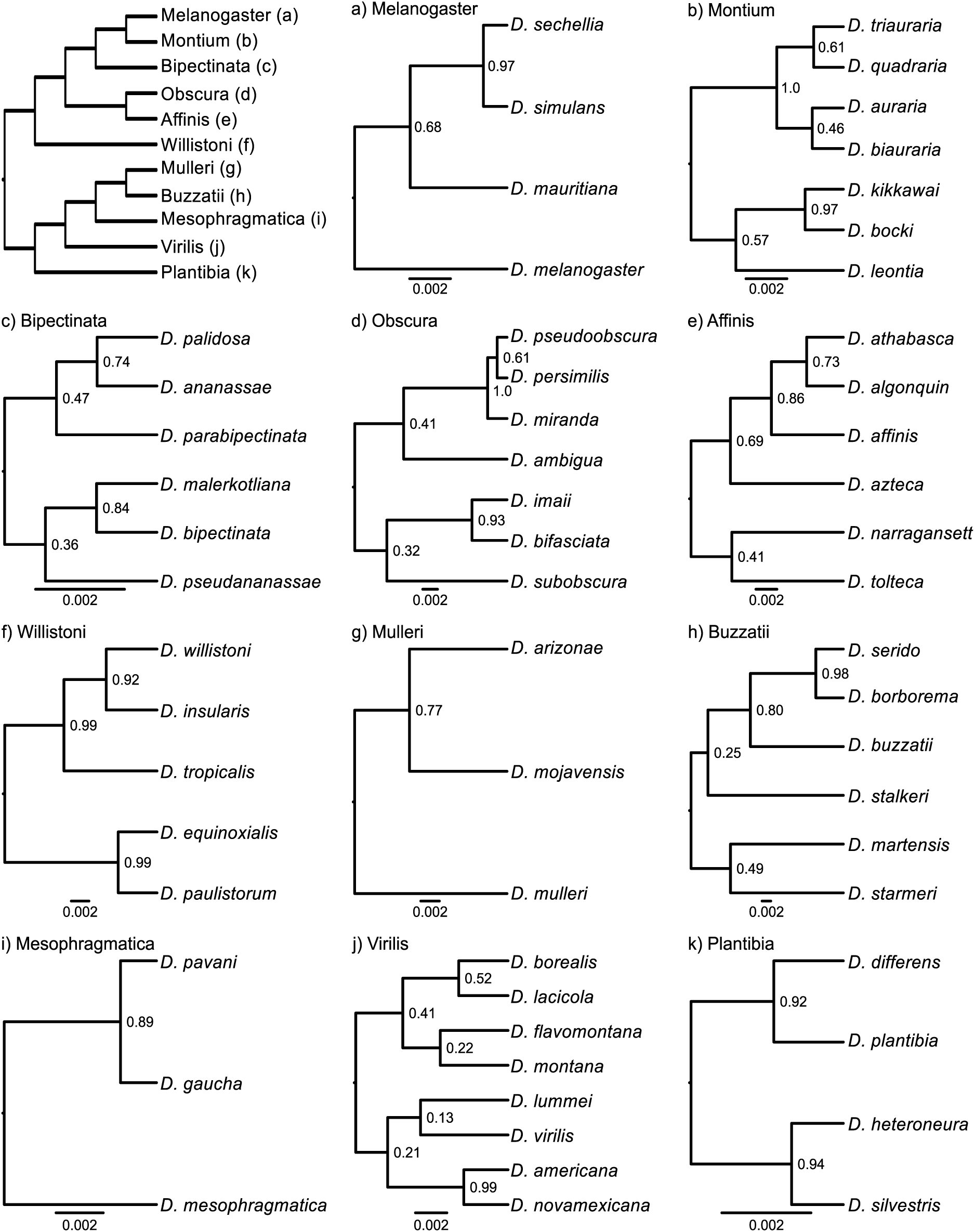
Consensus species trees for each species group in this study. Branch lengths (in coalescent units) are based on the average branch length from every MCMC tree that was topologically concordant with the highest probability species tree (taken from BPP output). Note that the scale bar is a different length for each tree because it represents the same value (0.002 coalescent units) for all trees. Backbone tree (top left) modified from Castillo (2017).

### Concordance of reproductively isolated species and coalescent species delimitation

The two approaches (i.e., threshold of genetic distance and reproductive isolation vs. PPIL threshold) recognized 63 out of the 99 species pairs as comprising two independent lineages. An additional 13 species pairs were not recognized as independent lineages by either approach. The two approaches were therefore in agreement on the species status for 76 of 99 species pairs in total (∼77%; Fig. 2). Of the remaining 23 species pairs, 17 met the threshold for independent lineages based on PPIL only, whereas the other six species pairs were considered to be distinct species based on total reproductive isolation (T) and genetic distance (D) only. Qualitatively similar, but somewhat more discordant, results were recovered when using the uninformed priors (68% agreement between methods). There was greater concordance between methods when delimiting sympatric taxa than allopatric taxa (Fig. 2) and this pattern held true across the informed and uninformed priors. When lowering the admittedly arbitrary, but common, PPIL threshold from 0.95 to 0.85 we recovered roughly the same amount of concordance (75 out of 99 species pairs in agreement; ∼76% concordance). However, there were more lineages recognized based on PPIL alone (Fig. S2). We also found greater discordance between methods for closely related species pairs (D ≤ 0.5) than those more distantly related (D > 0.5; Fig. S3).

**Figure 2).**
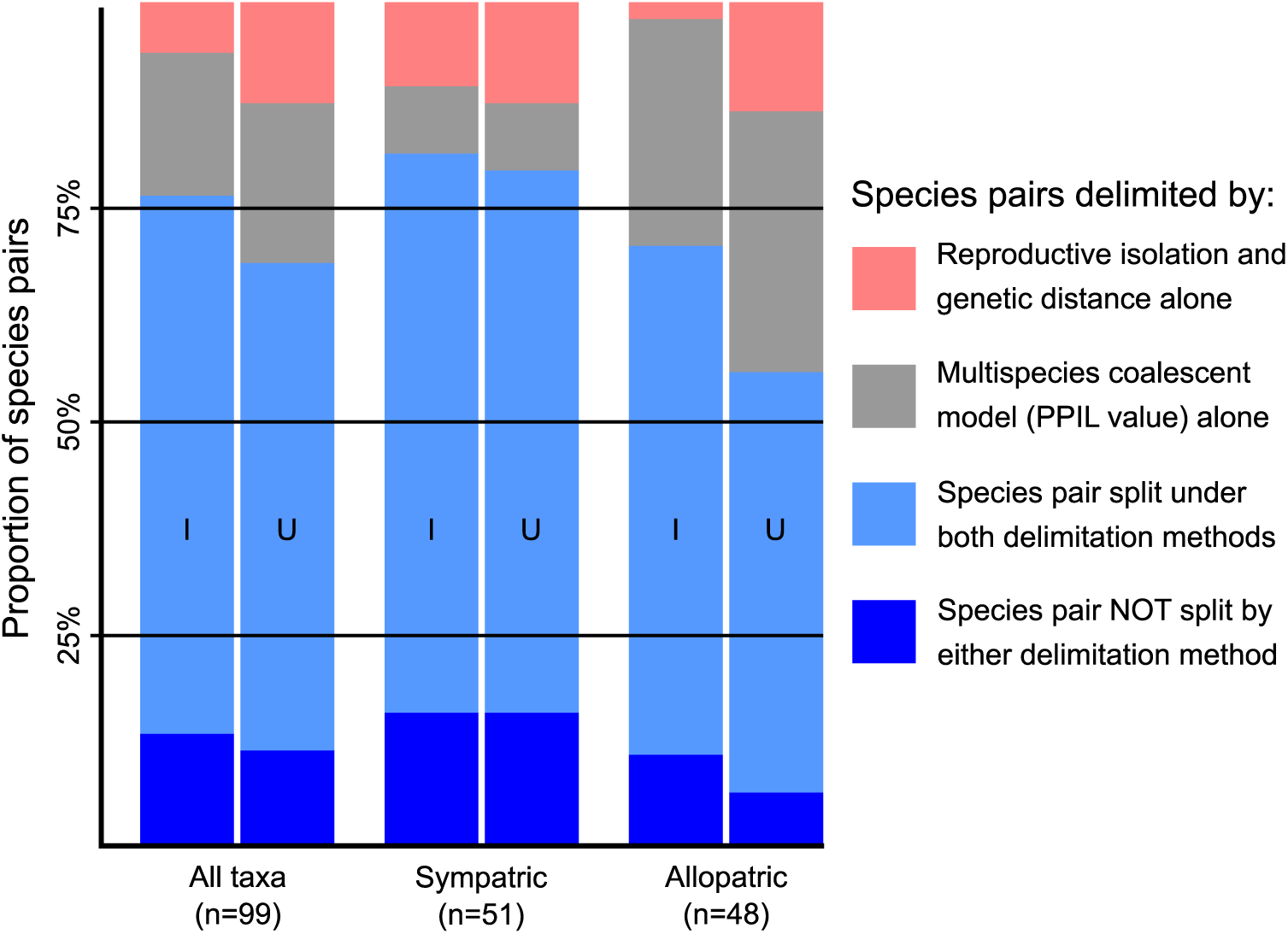
Plots comparing concordance between species delimitation models. Species delimitation models were in agreement for the majority of species pairs (Both; blue). However, there was discrepancy between certain species pairs. Some were split solely on reproductive isolation and genetic distance (RI + D; red) while others only on the basis of PPIL value ≥ 0.95 (MSC; gray). Results are shown for both informed (I) and uninformed (U) prior settings.

### Reproductive Isolation and Genetic Distance

We recovered better predictive models of PPIL when considering sympatric and allopatric taxa separately for prezygotic isolation and genetic distance, and similarly plausible models for postzygotic isolation (Table 1). When fitting the second set of GAMs to the different types of species pairs we found that the models of prezygotic isolation and genetic distance generally explained more null deviance than those for postzygotic isolation, which essentially reduced to a linear model (Fig. 3; plots for other GAM models are provided in Fig. S4 and Fig. S5). For allopatric taxa, the combined effect of prezygotic isolation + genetic distance was the best predictor of PPIL, but the effect of prezygotic isolation alone was similarly plausible (ΔAIC = 1.11; Table 2). In sympatric taxa, prezygotic isolation was the best predictor of PPIL, followed closely by the similarly plausible combination model of prezygotic isolation + genetic distance (ΔAIC = 0.21; Table 2). In the sympatric species pairs with low genetic distance (D < 0.5), we found that the best predictor of PPIL was again prezygotic isolation, with models of genetic distance and postzygotic isolation + genetic distance similarly plausible (ΔAIC = 0.66 & ΔAIC =0.58, respectively). We consistently found that the null deviance explained was low for the postzygotic isolation models (Table 2). These results remained qualitatively unchanged for the reduced phylogenetically independent datasets, with prezygotic isolation being the best predictor of PPIL in allopatry and sympatry. We present the results from the full dataset here, while complete results from the phylogenetically independent dataset can be found in the Supplementary Material (Table S4).

**Table 1).** Results from a series of GAMs to test if relationships are different between allopatric and sympatric species pairs, where *** means that model had the lowest AIC value and is therefore the most plausible model. The dagger (†) indicates a similarly plausible model (ΔAIC ≤ 2; Burnham et al. 2002). The proportion of null deviance explained by the model (“Dev. Exp.”) is also provided.

| Number of Smoothers | Prez. Isolation<br>(n = 100) |  | Post. Isolation<br>(n = 74) |  | Gen. Distance<br>(n = 108) |  |
| --- | --- | --- | --- | --- | --- | --- |
| | $\Delta AIC$ | Dev.<br>Exp. | $\Delta AIC$ | Dev.<br>Exp. | $\Delta AIC$ | Dev.<br>Exp. |
| One (Allo/Symp Together) | 8.15 | 0.29 | *** | 0.07 | 1.85<sup>†</sup> | 0.22 |
| Two (Allo/Symp Separate) | *** | 0.40 | 1.94<sup>†</sup> | 0.07 | *** | 0.27 |

**Table 2).** Table of ΔAIC values, where *** indicates the most plausible model (i.e., which explanatory variable best explains PPIL value). The dagger (†) indicates a similarly plausible model (ΔAIC ≤ 2; Burnham et al. 2002). The plus sign (+) between variables indicates a model of the additive effect of two predictors. The proportion of null deviance explained by the model (“Dev. Exp.”) is also provided.

| Explanatory Variable | Allopatric (n = 28) |  | Allopatric (D≤0.50; n = 15) |  | Sympatric (n = 42) |  | Sympatric (D≤0.50; n = 23) |  |
| --- | --- | --- | --- | --- | --- | --- | --- | --- |
|  | ΔAIC | Dev. Exp. | ΔAIC | Dev. Exp. | ΔAIC | Dev. Exp. | ΔAIC | Dev. Exp. |
| Prezygotic Isolation | 1.11<sup>†</sup> | 0.51 | *** | 0.61 | *** | 0.21 | *** | 0.39 |
| Postzygotic Isolation | 14.80 | 0.08 | 8.22 | 0.07 | 1.25<sup>†</sup> | 0.07 | 4.86 | 0.07 |
| Genetic Distance | 11.19 | 0.29 | 5.34 | 0.34 | 2.44 | 0.05 | 0.66<sup>†</sup> | 0.33 |
| Pre. Isol. + Gen. Dist. | *** | 0.57 | 4.52 | 0.40 | 0.21<sup>†</sup> | 0.23 | 2.23 | 0.33 |
| Post. Isol. + Gen. Dist. | 13.08 | 0.29 | 6.18 | 0.40 | 2.88 | 0.08 | 0.58<sup>†</sup> | 0.39 |

**Figure 3).**
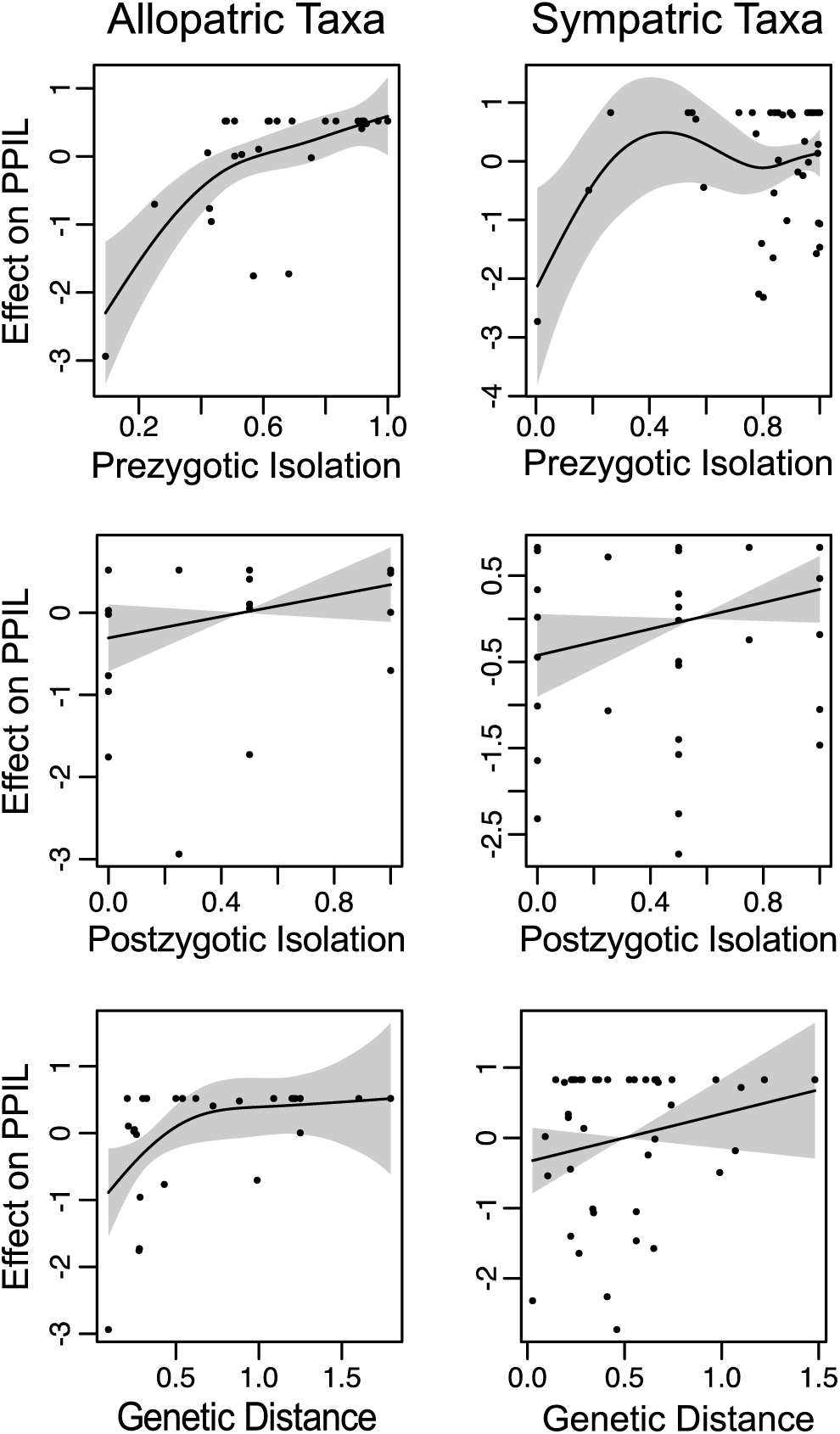
Fitted GAMs for allopatric (left) and sympatric (right) taxa. Note that the smooths are centered on zero to ensure model identifiability, meaning the y-axis is scaled relative to PPIL values but is not on the same scale (i.e., not from 0 – 1.0).

When looking at species pairs with prezygotic isolation data (n=100), we recovered a significant difference between the level of prezygotic isolation and PPIL value, particularly between species pairs with lower levels of prezygotic isolation (ANOVA; F-stat = 15.79; p-value = 2.01 × 10^−8^; Fig. 4a). However, when considering all species pairs with postzygotic isolation data (n=74), we found that there was no statistical difference in PPIL among the five levels of isolation (ANOVA; F-stat = 1.144; p-value = 0.343; Fig. 4b). Additionally, although the mean PPIL for species pairs that follow Haldane’s Rule was lower (0.927, n = 21) than those that do not (0.97; n = 11), there was no statistically significant difference in PPIL between the groups (t-test; p-value = 0.179; Fig. S6).

**Figure 4).**
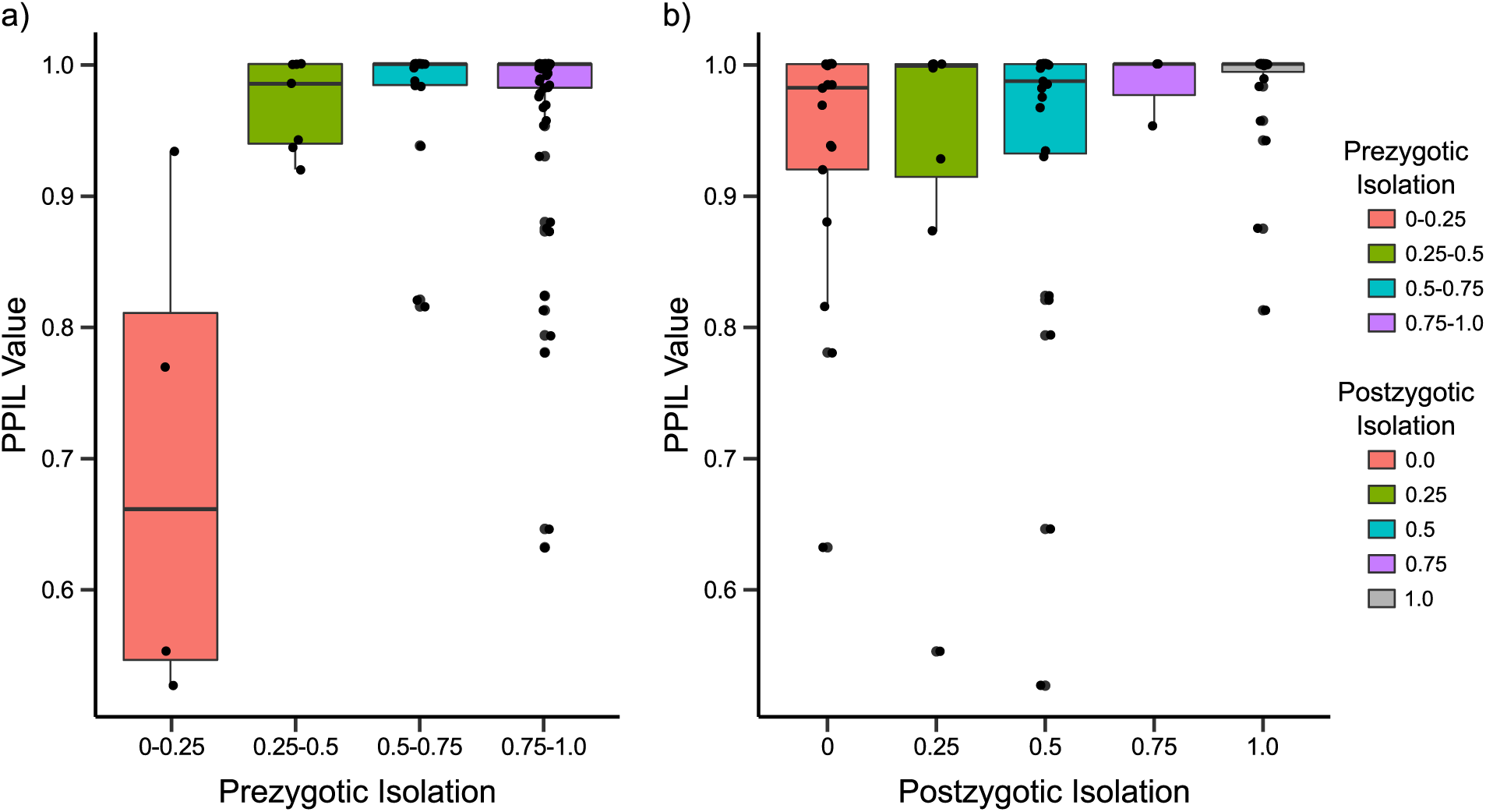
Boxplots depicting the results of ANOVA test across different levels of pre- or postzygotic isolation. There was a significant difference for prezygotic isolation (p-value = 2.01 × 10-8), but not for postzygotic isolation (p-value = 0.343).

### Effect of the Priors

We did not see an effect of the informed versus uninformed priors on the mean number of lineages delimited in a given group (i.e., BPP delimited a consistent number of species in each group across different prior settings). However, we did see a difference in the variance (i.e., the prior influenced how much uncertainty there was in the number of species in each group; Fig. 5b). Specifically, under the informed priors the MCMC samples were spread more evenly across possible delimitation models. We also observed a relatively large difference between the prior and posterior distributions of *Θ* and *τ* for analysis under uniformed priors, which was not observed in analyses under the informed prior (Fig. 5a). This indicates that the uninformed priors are a relatively poor match for *Drosophila*, which has the effect of biasing the results to be excessively certain. Moreover, this finding highlights that appropriate priors are important for accurately estimating the variance of a random variable, in addition to accurately estimating its mean.

**Figure 5a).**
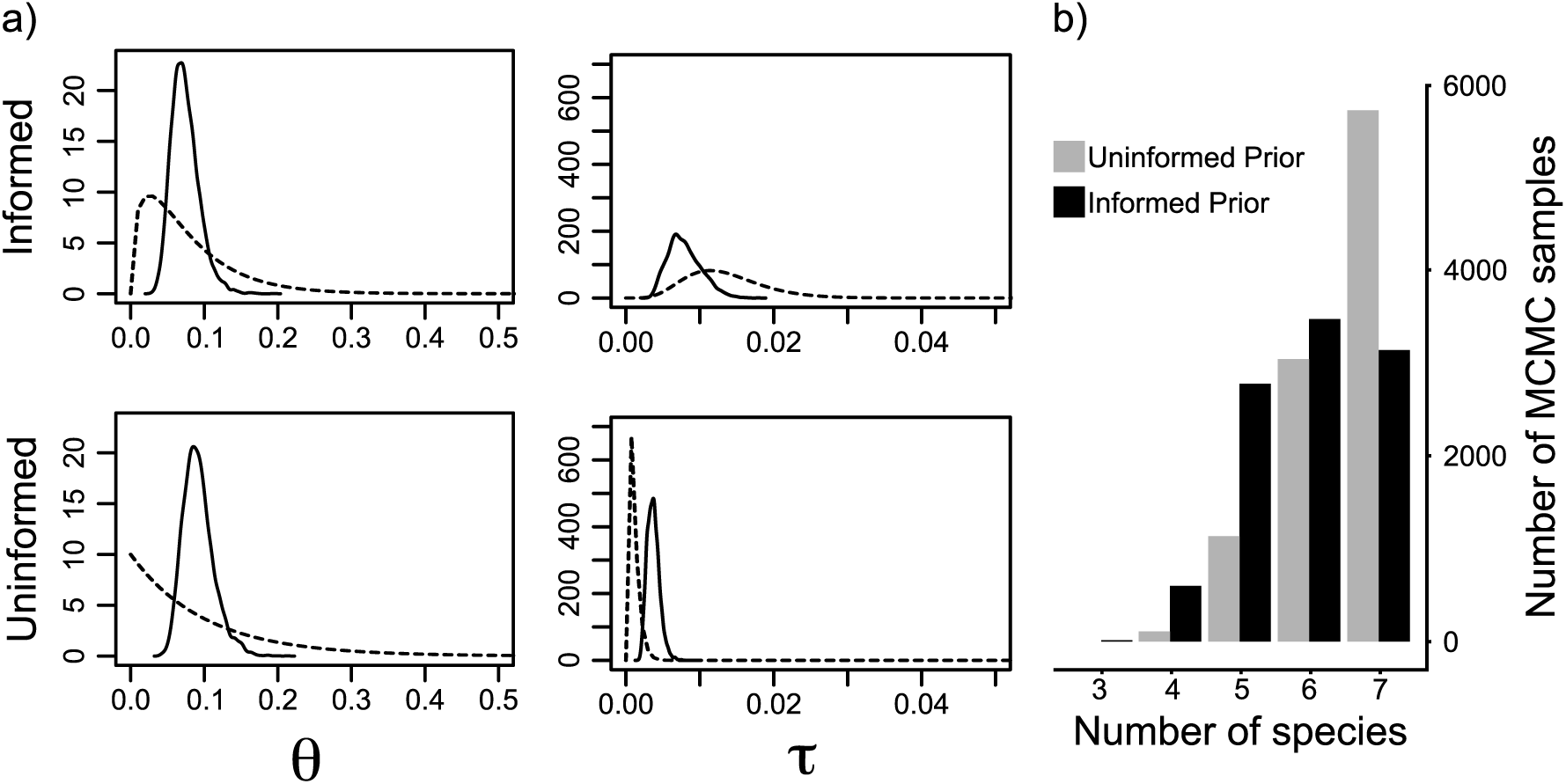
This figure compares relationships between prior (dashed) and posterior (solid) gamma distributions of *Θ* and *τ* using informed (top) and uninformed (bottom) prior distributions for the Montium group. Plots for all other species groups can be found in the Supplementary Material (Fig. S8). b) Shows the difference between the number of species delimited under unformed (gray) and uninformed (black) prior setting out of the total 10,000 MCMC samples per run.

## DISCUSSION

### Concordance of Species Boundaries Under the MSC

We assessed concordance between species that are delimited under population genetic models of lineage divergence with a biologically important and widely used measure of species distinctiveness, reproductive isolation. Specifically, we looked at species delimitations inferred under the BPP implementation of the MSC (which we consider an operational framework for a lineage-based species concept; Simpson 1951; Wiley 1978; De Queiroz 2007), and those considered distinct species determined by the amount of reproductive isolation and genetic distance between two lineages (i.e., a modified Biological Species Concept; Mayr 1963; Coyne and Orr 2004).

We find that the methods do not return identical results, and the multispecies coalescent tends to more readily split species than measures of reproductive isolation and genetic distance alone. This could result from the MSC recognizing genuine species that might be missed by other approaches. Specifically, our results confirm that (if we take MSC delimitation as truth) reproductive isolation does not need to be complete in order for lineages to be identifiable as independent. Alternatively, recent studies have found the MSC may be prone to oversplitting and might result in delimiting “population structure, not species” (Sukumaran and Knowles 2017; Chambers and Hillis 2019). However, because we are attempting to delimit nominal species any population structure within species should not mislead our interpretations of the species delimitation models. That being said, this also means that our results may provide a somewhat optimistic view of MSC performance.

When using the PPIL ≥ 0.95 cutoff, both criteria agreed with one another in the majority, but far from all, cases (76% agreement; PPIL values for all species pairs found in Table S5). Of these species pairs, 63 were delimited by both methods and 13 were delimited by neither. The cases where neither delimitation model recognizes these nominal taxa as independent lineages represent scenarios in which the taxonomy may require reexamination. For example, none of the species pairs in the Auraria species complex were recognized as independent by either method, suggesting current taxonomy in this species complex may be recognizing too many species.

Watada et al. (2011) recently took one step in this direction, revising the taxonomy of *D. quadraria* and suggesting it is better categorized as a junior synonym of *D. triauraria*.

Of the remaining species pairs (n = 23) for which there is disagreement between methods, the majority (n=17) were pairs split under the MSC (i.e., high PPIL) but not under the Coyne and Orr criteria. Most of these high PPIL pairs did not have enough total reproductive isolation (n = 14) to meet the Coyne and Orr threshold, while the remaining three species pairs did not have great enough genetic distance. These three species pairs are all allopatrically distributed. Due to their high levels of reproductive isolation (between 0.94-1.0), we expect that they would remain distinct upon secondary contact.

Of the 14 species pairs failing to meet the minimum total reproductive isolation threshold, 10 were allopatrically distributed. Because laboratory tests on reproductive isolation do not take geographic or environmental reproductive isolation into account, it is possible that allopatric pairs have not evolved a high degree of pre- or postzygotic isolation simply because they are genetically isolated by virtue of their distribution (i.e., geographic isolation is a strong barrier to gene flow, in itself). For example, the allopatric species pair of *D. americana* and *D. virilis* had relatively low genetic distance (D = 0.54) and total isolation (T = 0.644), but were never lumped under the MSC model (PPIL = 1.0). Furthermore, detailed crossing experiments and QTL mapping have shown high levels of postmating, prezygotic isolation between these two species (Sweigart 2010), suggesting that genetic distance and total reproductive isolation may take more time to evolve relative to isolation identified under a coalescent framework. The other four species pairs with high PPIL but low reproductive isolation were all sympatric. These may represent cases where BPP is identifying incipient species with (potentially) low levels of ongoing or recent gene flow (Leaché et al. 2019), or cases where the amount of reproductive isolation may have been underestimated. For example, the sympatric species pair *D. lummei – D. virilis* would be considered independent lineages under the MSC (PPIL = 1.0) despite having the lowest total isolation value in the entire dataset (T = 0.263). However, at least one source (Heikkinen and Lumme 1991) reported high postzygotic isolation between the pair that was not reflected in either Coyne & Orr (1989, 1997) or Yukilevich (2012), and therefore was not included our calculation of total isolation for this species pair.

The remaining six species pairs in disagreement would be considered species based on reproductive isolation and genetic distance, but were not identified as independent lineages under the MSC. At face value, this would indicate these are reproductively isolated species pairs that cannot be identified as independently coalescing lineages. Of these, five are sympatric, and only one is allopatric. For the only allopatric pair (*D. mesophragmatic – D. pavani*), the PPIL value was barely below the 0.95 posterior probability cutoff (PPIL = 0.942), and probably represents a valid split under both methods. We recovered a range of PPIL values for the five sympatric species pairs (PPIL between 0.794-0.930). Additionally, we observed that for all the sympatric species pairs with high reproductive isolation but PPIL ≤ 0.95 one or both species in that pair have documented chromosomal inversions (Jha and Rahman 1973; Johnson 1985; Noor et al. 2001).

In principle, any of these discrepancies could be attributed to error in measures of reproductive isolation, genetic distance, or PPIL (e.g., poor fit of the MSC to the data). Widespread error in reproductive isolation measurements seems unlikely due to the controlled laboratory conditions under which these data were collected. However, extrinsic isolating mechanisms (e.g., ecological selection against hybrids) are not considered in a laboratory environment, and may represent a greater source of discordance between laboratory and nature than anything else. The amount of genetic distance could also have been overestimated or underestimated, but this seems unlikely to be a systematic issue given that we found a strong correlation between Nei’s electrophoretic distance and Jukes-Cantor genetic distance based on COII (Fig. S1). While it could be true that species pairs are not as distinct as the amount of genetic distance and reproductive isolation between them might suggest, it is also possible that we failed to delimit them due to a lack of statistical power, driven by a paucity of loci used to fit the MSC model. The lowest PPIL among reproductively isolated species (*D. algonquin – D. athabasca*; PPIL = 0.79) was based on sequence data from only one individual per species for two genes, which is among the lowest amount of data for any species pair in the entire dataset (Table S3). Discordance between methods in these cases may simply be driven by the amount of data available for analysis, rather than a true disagreement between methods. However, performance in BPP is good even with a small number of loci (Rannala and Yang 2017), and we found no relationship between the number of loci compared between species pairs and the PPIL value for that species pair (linear regression: R^2^ = 0.002; p-value = 0.61; Fig. S7).

### Reproductive Isolation and GAMs

We observed significant model improvement when specifying separate smoothers for sympatric and allopatric taxa, rather than considering all taxa jointly, for most explanatory variables (Table 1). Based on these results, we subsequently explored the predictive power of three variables, and two combinations of reproductive isolation and genetic distance (see Materials & Methods), to predict PPIL values for sympatric and allopatric taxa separately. In allopatric taxa, we found that the combination model of prezygotic isolation + genetic distance best predicted PPIL value, with prezygotic isolation alone being found to be similarly plausible (ΔAIC = 1.11). This indicates that knowledge about prezygotic isolation, particularly in conjunction with genetic distance, is a strong predictor of PPIL values for allopatric species pairs. Although this makes sense at face value, it is interesting that prezygotic isolation represents a good predictor of speciation for taxa whose ranges do not overlap and therefore do not have the potential to mate in nature (i.e., prezygotic isolation does not matter if taxa are allopatrically distributed). Furthermore, the fact that the model of postzygotic isolation was the worst predictor of PPIL for allopatric species pairs (ΔAIC = 14.8) suggests the fate of hybrids between allopatric taxa, either through lab trials or secondary contact, says little about the independence of those lineages under the MSC. Models of genetic distance and postzygotic isolation + genetic distance were also poor predictors of PPIL in allopatric taxa. Taken together, this seems to imply that neither the time since two species became geographically isolated (inferred from genetic distance) nor levels of hybrid sterility/inviability are particularly indicative of lineage independence in allopatric taxa.

In sympatric taxa, we found that the model of prezygotic isolation alone was a better fit than all other explanatory variables, although the combination model of prezygotic isolation + genetic distance represented a similarly plausible alternative model (ΔAIC = 0.206). This agrees with the expectation that prezygotic isolation should be important in maintaining species boundaries in sympatry. When looking at sympatric taxa at low genetic distance (D < 0.5), we found the combined model of prezygotic isolation + genetic distance model best predicted PPIL. This suggests that time since divergence (inferred from genetic distance), and not just the amount of prezygotic isolation, is an important indicator of lineage independence among closely related and recently diverged sympatric taxa. The model for postzygotic isolation was the worst predictor of PPIL for closely related sympatric taxa (ΔAIC = 4.56). Furthermore, in contrast to the finding from Coyne and Orr (1989, 1997) we find that prezygotic isolation is the best predictor of PPIL in closely related (D ≤ 0.50) allopatric *and* sympatric species pairs. Given that reinforcement, by definition, does not occur in allopatry (i.e., there are no unfit hybrids), this reaffirms the relative importance of prezygotic isolation throughout the speciation process, even in the absence of any postzygotic isolation.

ANOVA tests revealed a significant difference between the level of prezygotic isolation and PPIL, but not between level of postzygotic isolation and PPIL, which recapitulates the results of the GAM analysis. Furthermore, this shows that once a low level of prezygotic isolation has been obtained (≥ 0.25) there is no statistical difference in PPIL values at higher levels of prezygotic isolation. Lastly, here was no statistically significant difference in PPIL for partially reproductively isolated species pairs (postzygotic isolation between 0.25-0.75) that do or do not show signature of Haldane’s Rule. Although there is abundant support for the occurrence of Haldane’s Rule in nature (Delph and Demuth 2016), sterility/infertility of the heterogametic sex does not seem to serve as a strong predictor of whether two lineages are identified as independent coalescent lineages.

### Effect of the Prior

Overall, we observed relatively consistent PPIL values across all prior settings (Table S4), indicating results from BPP analyses are not especially sensitive to the prior for either *Θ* or *τ*. As we would expect, we observed larger differences between the prior and posterior distribution of *Θ* and *τ* for the uninformed priors than the informed priors (Fig. 4a). The informed priors allowed the MCMC to more widely explore topological space, both in terms of the number of species in a given topology and the frequency of proposing a unique topology. Uniformed priors are likely to depart strongly from empirically reasonable values for any particular dataset, which can constrain how well the MCMC explores areas of parameters space that may have high likelihoods (but very low prior probability). While we did not observe strong impacts on the mean number of species that were inferred for these data, the misspecified priors did lead to inflated estimates of certainty (Fig. 4b). Given the existing concerns regarding whether the MSC tends to oversplit lineages, an exaggerated increase in precision, and presumably accuracy in certain situations, would almost certainly exacerbate any innate complications of MSC species delimitation (see also Leaché et al. 2019 for an alternative approach to better characterizing uncertainty in MSC species delimitation).

### Implications for the practice of species delimitation

Here we present a detailed look at the difference between two species delimitation approaches in a well-studied model organism, but our findings have implications beyond *Drosophila* speciation. First, the overall amount of discordance between the two delimitation methods we used here was much higher than we naïvely expected. If nominal taxa represent valid species, then any model of species delimitation should recover them. Moreover, while the frequency of speciation-collapse in nature is not yet well known, recent work has led to an increased recognition that the evolutionary history of many species is reticulate in nature (i.e., periods of gene flow during and/or after speciation; Burbrink and Gehara 2018; Marques et al. 2019) or otherwise non-bifurcating (e.g., budding or anagenic speciation; Silvestro et al. 2018). Taken together, our results confirm that (if we take MSC delimitation as truth) reproductive isolation does not need to be complete in order for lineages to be identifiable as independent.

Alternatively, the MSC model may not be able to pick up on reproductive isolating mechanisms that are specific to particular regions of the genome (i.e., genomic islands of divergence; Wolf and Ellegren 2016). In this study, we found that for sympatric species with high reproductive isolation, but low PPIL value, chromosomal rearrangement could be implicated in the maintenance of species boundaries. Moreover, while chromosomal rearrangements have long been associated with reproductive isolation and speciation (Rieseberg 2001; Campbell et al. 2018), to our knowledge have not been formally investigated under a MSC model. Supposing chromosomal inversions drive a rapid increase in reproductive isolation, the effects of localized genomic islands of divergence may not extend throughout the genome or be reflected in the few loci being used for delimitation. Although selecting a number of genes from a heterogeneous gene pool will often provide a useful approximation of species limits, if the only difference between species are relatively small genomic islands of divergence (e.g., chromosomal inversions) the results from MSC delimitation may be inconsistent with the levels of reproductive isolated observed in nature.

This study also allowed us to reveal novel insight into the speciation process. Most notably, we found that levels of prezygotic isolation were consistently related to PPIL, and found no indication that postzygotic isolation was similarly informative. Although quantification of pre- and postzygotic isolation are not readily available outside of model organisms, the implications of this finding potentially extends beyond the lab. Specifically, they may indicate that prezygotic isolation evolves earlier in the speciation process and rapidly leads to identifiable independent lineages, even under a relatively small amount of assortative mating (prezygotic isolation ≥ 0.25). This may provide some insight into why hybridization appears to be more common in nature than was previously appreciated (Taylor and Larson 2019). As long as most of the gene flow is within species, the boundaries are maintained and the speciation process in not disrupted. However, further work is needed to move beyond such speculation. Overall, our results highlight the value of considering the processes that underlie speciation, in conjunction with the assumptions of each species delimitation method employed, as a fruitful means for understanding conflicts (see also Barley et al. 2018; Smith and Carstens 2018).

## CONCLUSION

While reproductive isolation has long been a cornerstone of speciation research, it can rarely be used as an operational delimitation method because collecting such data is not tractable for most systems. Likewise, many practitioners of model-based species delimitation do not take reproductive isolation into account because the data are unavailable. In this study, we focus on a *Drosophila* dataset that is uniquely suited to address this divide in how species are understood. Here, we formally investigated how species boundaries based on reproductive isolation compare to model based species delimitation. We found that model based and reproductive isolation based methods agreed on ∼77% of species delimitations. MSC consistently delimited more species pairs (∼17%), while 6% of the species pairs were delimited based on reproductive isolation and genetic distance alone. However, because our study design does not allow for within species population structure to be misinterpreted as species boundaries, our results may provide a somewhat optimistic view of MSC performance. Additionally, by using a set of informed and uninformed prior settings we were able to qualitatively assess the accuracy and precision of MSC species delimitation, as implemented by BPP. In accordance with earlier studies, BPP appears to be relatively robust to prior misspecification, although we do find evidence that prior misspecification can lead to overconfidence in estimates of species boundaries. Lastly, based on predictive GAM models, we found that the amount of postzygotic isolation between species pairs was relatively uninformative as to whether or not those lineages can be recognized as independent under the coalescent. We are not questioning that postzygotic isolation acts as a strong reproductive barrier, certainly it does. However, it seems that prezygotic isolation evolving early in the speciation process is closely tied to recognizably independent lineages. As such, prezygotic isolation may be relatively more important in the incipient stages of speciation, and possibly to the process as a whole.

## SUPPLEMENTARY MATERIAL

Data available from the Dryad Digital Repository: XXXXXX

## FUNDING

This work was supported by the US National Science Foundation (grant numbers DEB 1354506, DEB 1754350, and DBI 1356796). This work was also supported by computational resources provided by the University of Hawai‘i Cyberinfrastructure group, as well as funding from the Arnold and Mabel Beckman Foundation.

